# A Novel Monoclonal Antibody Targeting a Large Surface of the Receptor Binding Motif Shows Pan-neutralizing SARS-CoV-2 Activity Including BQ.1.1 Variant

**DOI:** 10.1101/2023.01.20.524748

**Authors:** Leire de Campos-Mata, Benjamin Trinité, Andrea Modrego, Sonia Tejedor Vaquero, Edwards Pradenas, Natalia Rodrigo Melero, Diego Carlero, Silvia Marfil, Anna Pons-Grífols, María Teresa Bueno-Carrasco, César Santiago, Ferran Tarrés-Freixas, Victor Urrea, Nuria Izquierdo, Eva Riveira-Muñoz, Ester Ballana, Mónica Pérez, Júlia Vergara-Alert, Joaquim Segalés, Carlo Carolis, Rocío Arranz, Julià Blanco, Giuliana Magri

## Abstract

In the present study we report the functional and structural characterization of 17T2, a new highly potent pan-neutralizing SARS-CoV-2 human monoclonal antibody (mAb) isolated from a convalescent COVID-19 individual infected during the first wave of the COVID-19 pandemic. 17T2 is a class 1 VH1-58/κ3-20 antibody, derived from a receptor binding domain (RBD)-specific IgA memory B cell and developed as a human recombinant IgG1. Functional characterization revealed that 17T2 mAb has a high and exceptionally broad neutralizing activity against all SARS-CoV-2 spike variants tested, including BQ.1.1. Moreover, 17T2 mAb has *in vivo* prophylactic activity against Omicron BA.1.1 infection in K18-hACE2 transgenic mice. 3D reconstruction from cryogenic-electron microscopy (cryo-EM) showed that 17T2 binds the Omicron BA.1 spike protein with the RBD domains in “up” position and recognizes an epitope overlapping with the receptor binding motif, as it is the case for other structurally similar neutralizing mAbs, including S2E12. Yet, unlike S2E12, 17T2 retained its high neutralizing activity against all Omicron sublineages tested, probably due to a larger contact area with the RBD, which could confer a higher resilience to spike mutations. These results highlight the impact of small structural antibody changes on neutralizing performance and identify 17T2 mAb as a potential candidate for future therapeutic and prophylactic interventions.

## Introduction

SARS-CoV-2, the etiological agent of COVID-19, has provoked one of the worst pandemics in human history, causing more than 6.7 million deaths registered worldwide (https://covid19.who.int/). The high level of virus circulation amongst humans and other species has led to the emergence of several variants of concern (VOCs) with progressively increased transmissibility and immune evasion capacity ^1–3^. In December 2021, the Omicron VOC became the globally dominant circulating strain, after replacing previous variants. The initial Omicron wave was caused by the BA.1 variant, followed by several Omicron sublineages (BA.2, BA.5 and BQ.1.1 among others). Compared with the ancestral variant identified in Wuhan (WH1), the spike protein of the BA.1 one contains at least 30 amino acid (aa) substitutions including 6 aa deletions and 3 aa insertions which are largely confined to the receptor binding domain (RBD) and the N-terminal domain (NTD), the two major antigenic sites targeted by neutralizing antibody responses ^4^. BA.2 shares 21 mutations with BA.1 and harbors an additional 8 specific mutations. BA.5 appears to have evolved from BA.2 with two additional mutations and one reversion in the RBD and one deletion in the NTD ^1^. Since November 2022, BQ.1 and BQ.1.1 variants, which harbor 3 new mutations in key antigenic sites of the RBD, have overtaken BA.5 as the dominant variant across the USA and Europe ^5^. The presence of these mutations has not only increased the transmissibility of Omicron variants but has also caused a strong resistance to vaccine-induced antibody responses, leading to unprecedented levels of vaccine breakthrough infections worldwide^1,6,7^.

Together with vaccines and antiviral drugs, neutralizing monoclonal antibodies (mAbs) targeting the spike protein of SARS-CoV-2 have been extensively used to treat individuals at the highest risk of severe COVID-19 ^8,9^. During previous COVID-19 waves, therapeutic administration of mAbs was reported to be highly effective in preventing COVID-19-related hospitalization and death ^10,11^. However, most of the mAbs currently in use were developed against ancestral SARS-CoV-2 and they all lost or significantly reduced their activity against highly mutated Omicron sublineages ^1,2,12^. Therefore, there is an urgent need to develop new pan-SARS-CoV-2 neutralizing antibodies that are effective against current and future VOCS to provide additional therapeutic options to patients.

Here we report the functional and structural characterization of a new human monoclonal antibody developed from a patient infected with the ancestral SARS-CoV-2 that shows pan-neutralizing activity against both pre-Omicron and Omicron SARS-CoV-2 variants, including BQ.1.1.

## Results

### Generation of human recombinant SARS-CoV-2-specific monoclonal antibodies

To generate human mAbs capable of neutralizing SARS-CoV-2, we sorted 380 circulating RBD^+^ B cells from a convalescent COVID-19 individual who was infected during the first wave of the pandemic in Spain ^13^, using a biotinylated RBD protein from the ancestral SARS-CoV-2 strain as bait (**Supplementary Figure 1A**). RBD-specific B cells were further characterized according to the expression of CD27, CD21, CD11c, HLA-DR, IgM, IgD, IgA and the Ig light chain λ (**Supplementary Figure 1B**). mRNA from sorted cells were then reverse-transcribed and the immunoglobulin heavy chain (IGHV) and light chain variable (IGLV) regions were amplified by PCR following an established protocol ^14^. After Sanger sequencing, 5 RBD-specific IGHV and IGLV paired regions were cloned into expression vectors and generated as recombinant human monoclonal IgG1. Three of these mAbs were generated starting from RBD-specific CD21^+^CD27^+^ canonical IgG memory B cells whereas two of them, 17T2 and 54T1, were generated starting from RBD-specific IgA canonical memory B cells (**Supplementary Table 1**). As expected, all 5 mAbs showed relatively low levels of somatic mutations, which is consistent with the low level of hypermutation reported in RBD-specific antibodies following infection with the ancestral SARS-CoV-2 ^15^ (**Supplementary Table 1**). These mAbs were first screened to confirm their reactivity profile by enzyme-linked immunosorbent Assay (ELISA) using recombinant RBD from the WH1 and the subsequent Beta (B.1.351), Gamma (P.1), Delta (B.1.617.2), Omicron BA.1 and Omicron BA.2 variants as immobilized proteins. As expected, all mAbs bound to the RBD from the ancestral variant, yet only 17T2 was able to efficiently recognize all variants tested, including RBD from highly mutated Omicron variants (**Supplementary Figure 2**). The binding affinity of 17T2 for the RBD from different SARS-CoV-2 variants was then assessed by surface plasmon resonance (SPR). This analysis confirmed the high-affinity binding with equilibrium dissociation constants (K_D_) in the subnanomolar range for all variants tested (**Supplementary Table 2)**. Interestingly, 17T2 belongs to the IGHV1-58/κ3-20 clonotype, as many other potent neutralizing mAbs isolated from SARS-CoV-2-infected and/or vaccinated individuals ^16–21^ (**Supplementary Table 1**).

### 17T2 mAb shows high neutralization activity against all SARS-CoV-2 variants including Omicron sublineages

To evaluate the functional activity of the selected antibodies, we tested their neutralization capacity using HIV reporter pseudoviruses expressing different SARS-CoV-2 spike proteins from variants ranging from WH1 to Omicron BA.1 (**Supplementary Table 3**) ^22^. Consistent with the reactivity data, all antibodies showed neutralization of WH1 and D614G pseudoviruses. However, only 17T2 mAb maintained high neutralizing capacity against all variants tested, while the other mAbs markedly lost potency against Beta, Gamma, Mu or Omicron BA.1 variants **(Figures 1A** and **1B**). 131T2 showed the lowest potency against pre-Omicron variants and no activity against Omicron BA.1 (**Figure 1A**). To further characterize 17T2 mAb we analyzed neutralization of pseudoviruses exposing the spike of newer Omicron subvariants (**Supplementary Table 3**) and SARS-CoV-1. Moreover, for comparative purposes, we assayed in parallel two well-characterized broad-spectrum neutralizing RBD-targeting mAbs: S2E12 and S309 ^21,23^. S2E12 is a VH1-58/κ3-20-encoded class 1 antibody, like 17T2, whose antibody-binding epitope overlaps with the receptor binding motif (RBM) in the RBD ^20^, whereas S309 is a class 3 antibody isolated from a patient recovering from SARS, which binds to a conserved epitope outside the RBM ^24^. 17T2 mAb neutralized all SARS-CoV-2 variants tested (pre-Omicron variants WH1, D614G, Alpha, Beta, Gamma, Delta, and Omicron subvariants BA.1, BA.2, BA.4/5 and BQ.1.1) with IC50 values ranging from 38 to 2 ng/ml for WH1 and BA4/5, respectively. In contrast, no activity against SARS-CoV-1 was observed (**Figures 1C** and **1D**). The wide SARS-CoV-2 neutralization spectrum of 17T2 was remarkably different from the other two broadly neutralizing mAbs. S2E12 showed higher potency against all pre-Omicron variants but had significant lower neutralizing activity against BA.1, BA.2 and no activity against BA.4/5 and BQ.1.1 (**Figures 1C** and **1D)**, while the pan-neutralizing antibody S309 maintained some activity against Omicron subvariants, but with a significantly lower potency compared to 17T2 (**Figures 1C-E**).

**Figure 1.**
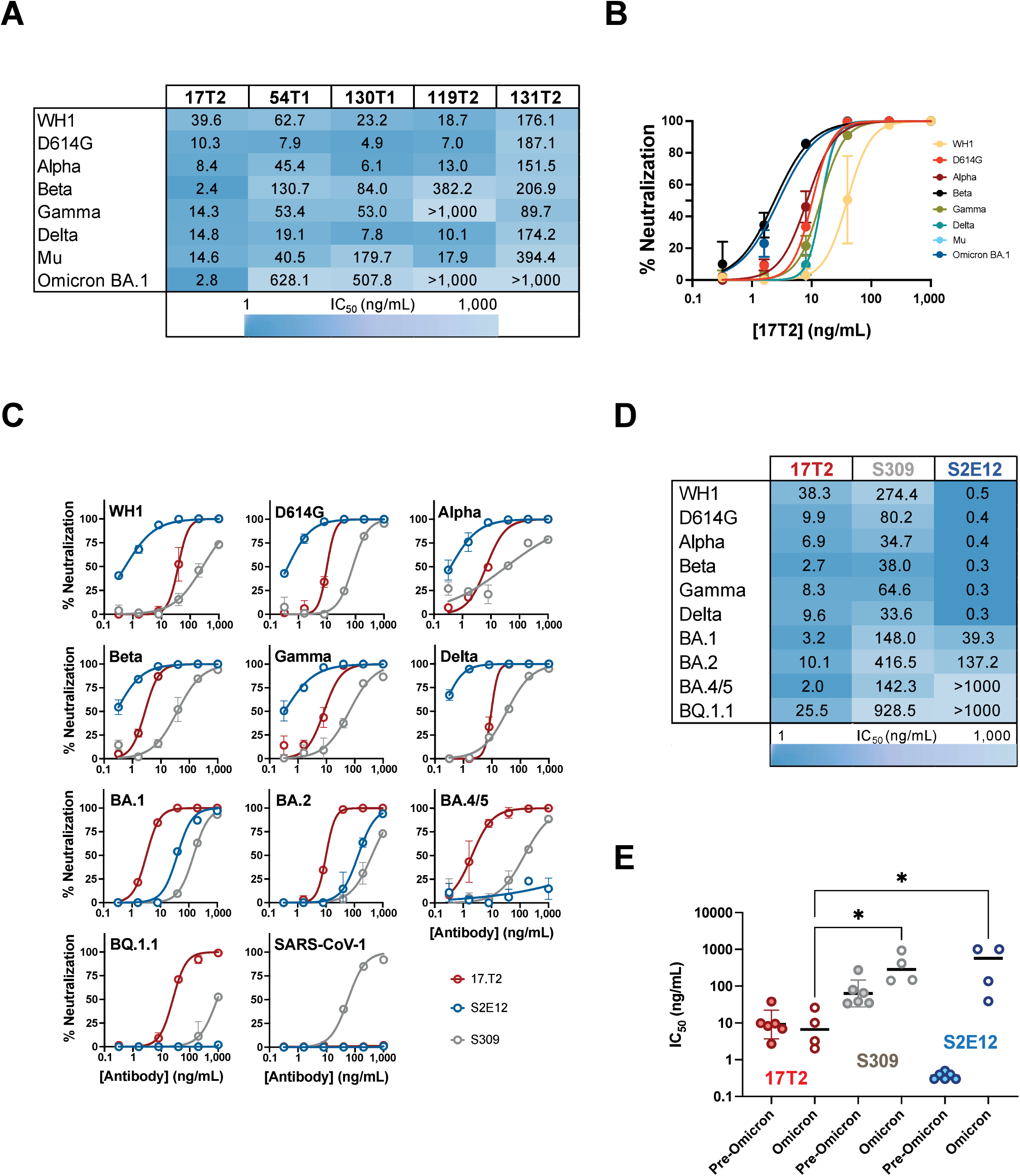
Pan-neutralizing activity of 17T2 mAb. (**A**). Neutralizing activities of the selected mAbs against the indicated SARS-CoV-2 variants. Values are in ng/mL and color coded as indicated in the bottom of the figure. (**B**) Neutralization curves of 17T2 mAb against the pseudoviruses tested in panel A. Values are mean ± SD of duplicate samples. (**C**) Neutralization curves of 17T2 (red), S309 (grey) and 2E12 (blue) mAbs against the indicated SARS-CoV-2 variants or SARS-CoV-1. Values are mean ± SD of duplicate samples (**D**) Summary of IC50 values from panel C in ng/mL and color coded as indicated in the bottom of the figure. (**E**) Impact of Omicron subvariants on neutralization capacity. IC50 values for pre-Omicron variants (WH1, D614G, Alpha, Beta Gamma and Delta) were grouped and compared to Omicron subvariants (BA.1, BA.2, BA.4/5 and BQ.1.1). Only comparisons of Omicron variants for the indicated mAbs are shown * (p<0.05, Kruskal-Wallis test with correction for multiple comparisons).

### 17T2 mAb has prophylactic activity against Omicron BA.1.1 *in vivo* in K18-hACE2 transgenic mice

Next, we tested 17T2 *in vivo* prophylactic efficacy in K18-hACE2 transgenic mice. 17T2 mAb or an isotype control antibody (IgGb12) were administered intraperitoneally (10 mg/kg) 24 hours before challenge with 10^3^ TCID50 of a BA1.1 SARS-CoV-2 isolate (**Figure 2A**). As previously reported ^25,26^, Omicron infection of K18-hACE2 transgenic mice resulted in a mild disease without significant changes in weight in infected animals as compared to uninfected ones (**Figure 2B**). However, isotype control-treated animals showed high viral loads in oropharyngeal swabs, lungs and nasal turbinates, both 3- and 7-days post infection (dpi) (**Figure 2C**). In comparison, mice treated with 17T2 mAb showed significantly lower viral loads in lungs 3 dpi and in all tissues assayed (oropharyngeal swabs, lungs, and nasal turbinate) at 7 dpi (**Figure 2C**). The protective effect of 17T2 was confirmed in lung tissue by analyzing histopathological lesions and the SARS-CoV-2 nucleocapsid focal expression at 3 and 7 dpi by histology and immunohistochemistry, respectively (**Figures 2D** and **2E**).

**Figure 2.**
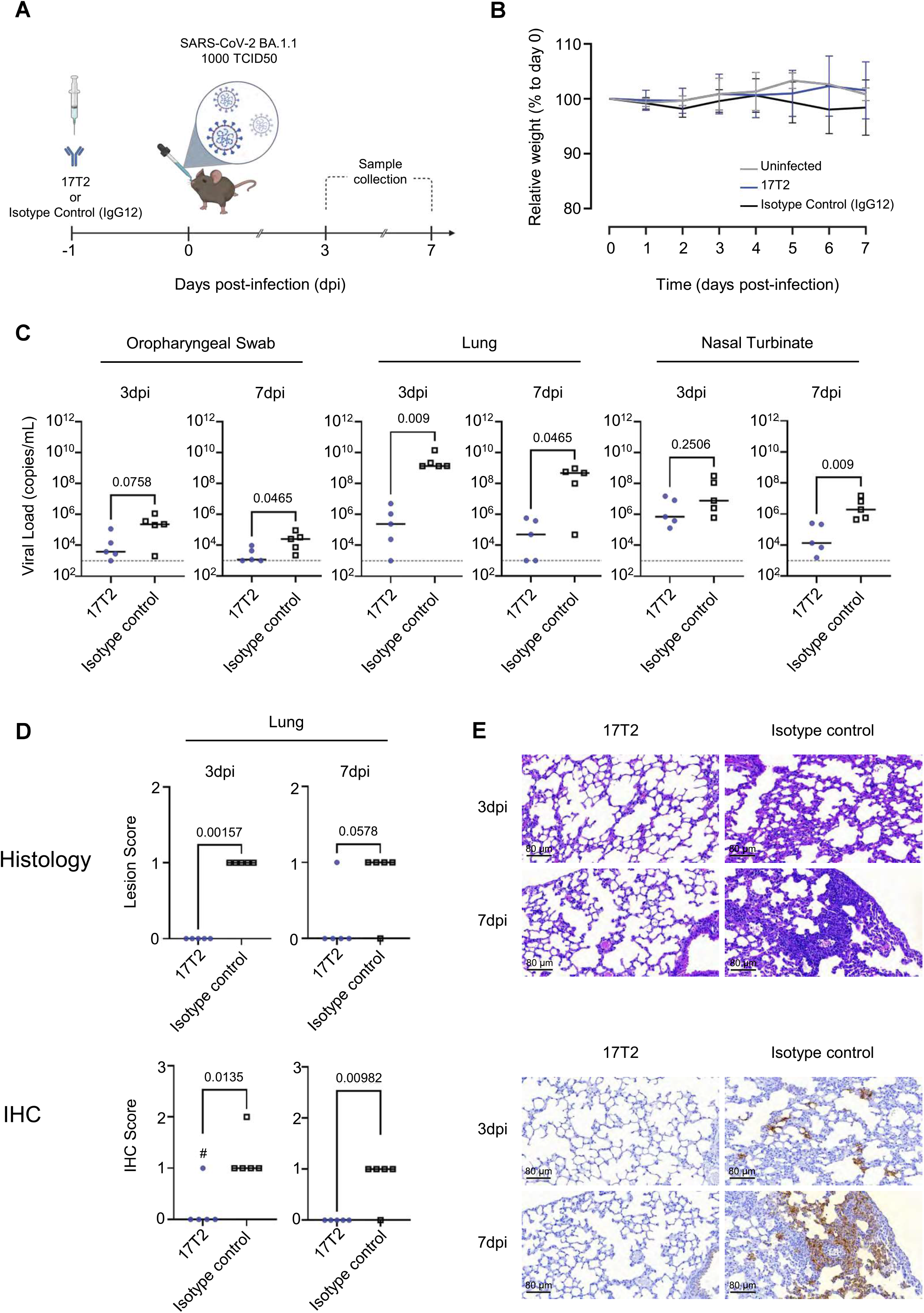
Prophylactic 17T2 protection against SARS-CoV-2 BA1.1 infection in K18-hACE2 transgenic mice. (**A**) Schematic description of the experimental setting. Transgenic K18-hACE2 mice were administered intraperitoneally with either 17T2 (17T2 mAb, n=10) or an isotype control (IgGb12, n=10). After 24h, treated animals were intranasally challenged with an Omicron BA.1.1 SARS-CoV-2 isolate (n=20), or PBS (Uninfected Control Group) (n=4). Mice were monitored for 7 days. Euthanasia was performed 3- and 7-days post-infection (dpi) (n=5 for each treated group per timepoint, n=2 uninfected per timepoint) for sample and tissue collection. Created with Biorender.com. (**B**) Relative K18-hACE2 transgenic mice weight-loss follow-up. Groups include: uninfected (grey, n=4), 17T2 mAb-treated and infected (blue, n=10), and isotype control-treated and infected (black, n=10) animals. Solid lines and bars represent mean± SD. (**C**) SARS-CoV-2 viral RNA load quantification (copies/mL) on different tissues: oropharyngeal swab, lung and nasal turbinate at 3 and 7 dpi in 17T2 mAb-treated (blue dots, n=5 per timepoint), isotype control-treated (black squares, n=5 per time point) infected animals. Limit of detection is represented by a dashed line. Statistically differences were determined using a Peto-Prentice generalized Wilcoxon test. (**D**) Histopathological and immunohistochemical scores of lungs from infected K18-hACE2 mice. Lesion (broncho-interstitial pneumonia) scoring: 0 = no lesion, 1 = mild lesion, 2 = moderate lesion, 3 = severe lesion. IHC scoring: 0 = no antigen, 1 = low and multifocal antigen, 2 = moderate and multifocal antigen, 3 = high and diffuse antigen. # indicates an animal which presented focal expression only visible on 1 out of 5 lung sections: it was scored as 1 but with minimal detection of virus replication. All other positive scores showed multifocal distribution on multiple lung sections. Comparisons were performed using an Asymptotic Generalized Pearson Chi-Squared Test for ordinal data with pairwise comparisons. (**E**) Representative histopathological and immunohistochemical findings, at 3 and 7 dpi, in K18-hACE2 transgenic infected mice treated with either 17T2 mAb or an isotype control. Histological slides were stained with hematoxylin and eosin, and immunohistochemistry ones were counterstained with Hematoxylin. Scale = 80 μM.

### 17T2 binds the Omicron BA.1 spike protein with the RBD domains in the up position and recognizes a large surface overlapping with the receptor binding motif

We carried out cryo-EM analysis to understand the potency and breadth of 17T2-mediated neutralization of SARS-CoV-2 variants. For this reason, we solved the structure of the complex between the highly mutated Omicron BA.1 trimeric spike and the 17T2 Fab fragment, reaching a resolution of 3.46 Å (**Figures 3A** and **3B**). Our analysis showed that 17T2 Fab binds to RBD in the “up” conformation with all particles containing 3 17T2 Fabs, each one bound to adjacent RBDs within a single spike trimer (**Figures 3A** and **3B**). Due to the higher conformational dynamics in the 17T2 variable domains and the RBD regions, resolution in the contact area was lower than in the rest of the trimer. Local refinement was performed in this region, significantly improving local resolution to 4.41 Å (**Figures 3C** and **Supplementary Figure 3**). After refinement, we determined that 17T2 Fab binds to the left shoulder-neck region of the RBD that is only accessible in “up” conformation (**Figures 3B-D**). The interaction site overlaps with the RBD in a similar manner to other class 1 VH1-58/κ3-20--derived neutralizing mAbs ^16–21^. Fab 17T2/RBD interactions involve both the heavy chain (HC) and light chain (LC) of the antibody, covering 563Å^2^ for the HC and 295 Å^2^ for the LC of the total interaction surface (**Figure 3D**). 17T2 Fab uses complementary determinant regions (CDR) 1 to 3 of the HC and CDR1 and CDR3 of the LC to recognize residues 420 to 421; 455; 473 to 478; 480; 484 to 487, 489 and 493 of the SARS-CoV-2 RBD (**Figures 3D** and **3E**). The contact area mostly involved Van der Waals interactions with a minor contribution from hydrogen bridges (**Figure 3E**). In addition, we identified a salt bridge formed between the D420 residue of the RBD and the R103 residue of H3 that stabilizes the binding of 17T2 to the RBD (**Figure 3E**). As previously described for structurally similar neutralizing antibodies ^16,27^, 17T2 is glycosylated at the N102 residue of H3 located near the left shoulder of the RBD (**Figure 3C**). Using PNGase-mediated de-glycosylation (**Supplementary Figure 4A**) we confirmed that this glycosylation had no effect on the affinity nor the neutralizing activity of 17T2 mAb (**Supplementary Figure 4B-D**).

**Figure 3.**
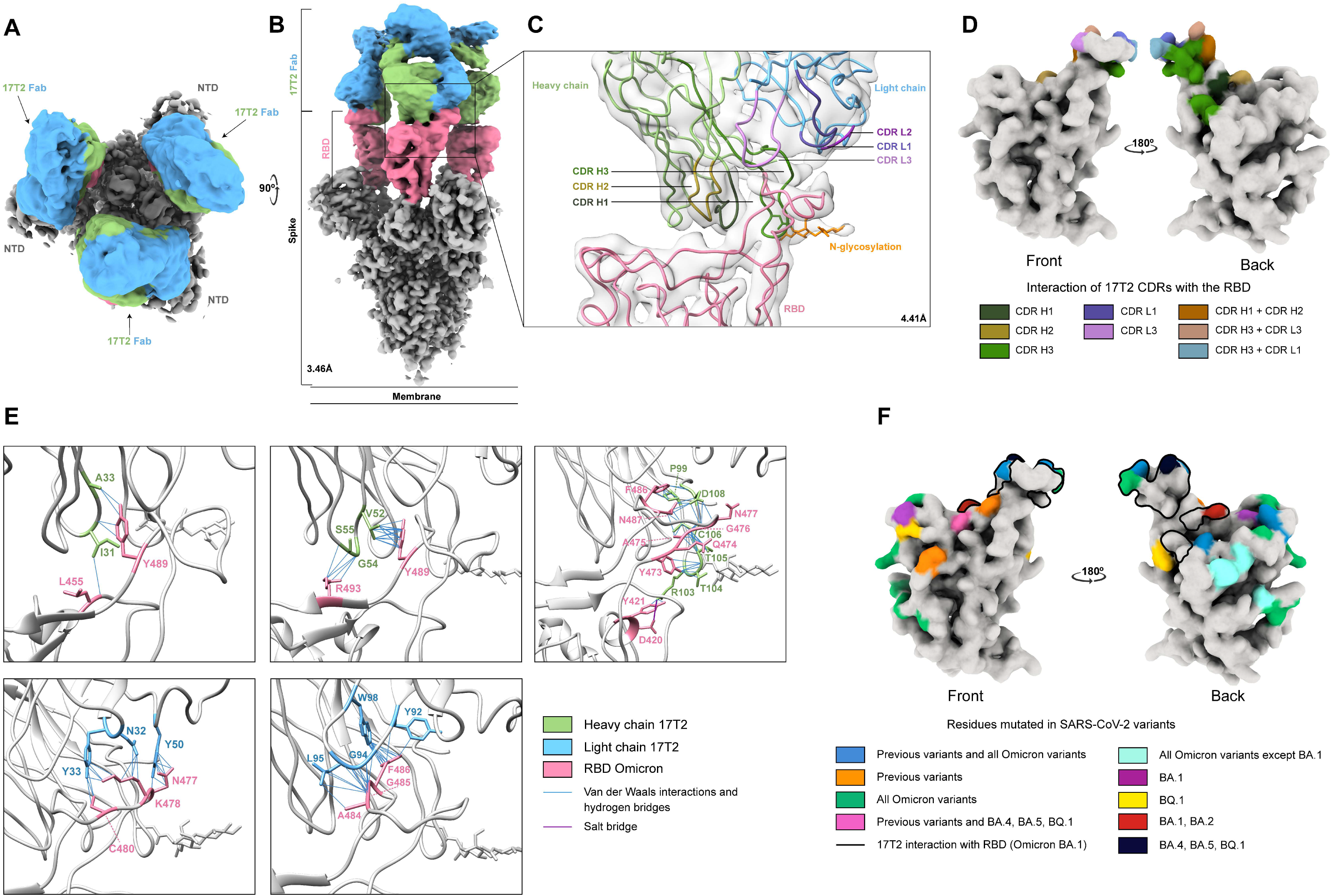
Structural and functional characterization of complex 17T2 Fab fragment with Omicron BA.1 spike using cryo-EM. (**A** and **B**) Top and side views of the cryo-EM map of Omicron BA.1 spike trimer with three 17T2 Fab fragments bound to three open RBDs. The core of the spike is shown in grey, the RBDs in pink, and the heavy chain (HC) and light chain (LC) 17T2 of the Fab in green and blue, respectively. (**C**) Structure of the RBD and 17T2 Fab after local refinement. The interaction zone between the RBD and the Fab is shown in cartoon representation where the three CDRs from each chain are distinctively colored and the N-glycosylation is indicated in orange. (**D**) CDRs that are involved in binding with RBD, specifically, interacting in the region of its left shoulder and neck (front and back view, respectively). (**E**) A detailed view of some interactions between 17T2 Fab and RBD. The main chains are colored in grey and the side chains of the residues involved in the interaction are shown in green for HC, in blue for LC, and in pink for RBD. (**F**) Locations of SARS-CoV-2 variant mutations on RBD relative to 17T2 epitope site that is shown as a black line (front and back view, respectively). The information about the variants and mutated residues can be found in **Supplementary Table 4**.

The 17T2 Fab binds to a large and mostly conserved area of RBD, with mutated residues harbored by SARS-CoV-2 variants located at the edge of the contact surface (**Figure 3F** and **Supplementary Table 4**). Since 17T2 and S2E12 mAbs share high sequence identity but differ in their neutralizing activity against highly mutated Omicron subvariants, we compared the structures of their respective Fabs/spike complexes (**Supplementary Figures 5A** and **5B**). This comparison revealed that S2E12 ^16,20,27^ and 17T2 Fabs bind parallel to the longest axis of the hACE2 binding site. Nevertheless, the area of interaction between 17T2 and RBD is broader than the area between S2E12 and RBD, the latter being included in the 17T2 interaction area (**Supplementary Figure 5C**). Moreover, although the structure of the two antibodies is highly similar, the amino acid side chains of 17T2 are located closer to the surface of the RBD (**Supplementary Figure 5B**), allowing for higher number of contacts with conserved residues. This fact probably contributes to the ability of this antibody to neutralize Omicron subvariants exposing S477N and F486V mutations, which strongly impact S2E12 binding.

## Discussion

Here we describe the functional and structural characterization of 17T2, a new human mAb with broad neutralizing activity against all SARS-CoV-2 variants, including Omicron sublineages BA.1, BA.2, BA.4, BA.5 and BQ.1.1. Importantly, prophylactic administration of 17T2 mAb resulted in significant reduction of viral replication and microscopic lung lesions in a murine model of SARS-CoV-2 Omicron infection.

17T2 belongs to the class 1 VH1-58/κ3-20-derived antibodies that include several mAbs (i.e. S2E12, Cv2.1169, A23.58.1, AZD8895) with high neutralizing activity against pre-Omicron variants ^16–18,20,21^. Yet, compared to these structurally similar mAbs, 17T2 retains its potency against all Omicron sublineages tested, including the highly immune evasive variants BA.5 and BQ.1.1 ^1,28^, with IC50 values ranging from 2 to 25.5ng/ml. The structural analysis of 17T2 Fab in complex with Omicron BA.1 spike trimer suggests complementary mechanisms to explain its broad neutralizing activity. On the one hand, the high antibody affinity allows for a complete blockade of all RBDs of the spike trimer stabilized in the “up” conformation (stoichiometry 3 Fab:1 spike trimer). On the other hand, when compared to S2E12, 17T2 shows a larger area of interaction with the RBM, which could confer higher tolerability to RBD mutations. Moreover, the presence of a salt bridge between the R103 in the CDR H3 and the D420 in a conserved region of the RBD participates in the stabilization of the complex, contributing to the extraordinarily high affinity of 17T2 to the RBD from multiple SARS-CoV-2 variants. Interestingly, D420 has been recently identified by mutagenesis as a potential site for escaping neutralization by some class 1 antibodies ^29,30^. The stability provided by D420 could explain why 17T2 mAb resists the F486V spike mutation present in the BA.4 and BA.5 variants which otherwise escape all other IGHV1-58-derived antibodies described thus far ^1,17,31^.

IgG Fab glycosylation appears to be a key parameter in immunity with possible consequences on antigen binding and antibody activity ^32^. Our structural analysis revealed that 17T2 is glycosylated at residue N102 in the CDR H3 region, in close proximity to the area of contact with the RBD. However, PNGase mediated de-glycosylation of 17T2 mAb had no effect on the binding to RBD or on the neutralizing activity, excluding a potential role of Fab glycosylation in its functional activity. Yet, we cannot rule out its possible implication in the stability or immune modulatory properties of the antibody^32^.

Our data indicates that 17T2 shows unique structural properties conferred by single point mutations in CDRs from both heavy and light chains (as compared to S2E12), enabling its potent and broad neutralizing activity. However, both the RBD exposure in the spike trimer and the presence of RBD mutations in the outer 17T2 mAb binding area seem to induce minor variations in the neutralization potency against the tested variants (IC50 range 3-38 ng/mL). A four-fold change in IC50 was observed when comparing WH1 and D614G pseudoviruses, which share identical RBD sequences but differ in the RBD “up” and “down” conformational equilibrium. Similarly, mutations such as S477N, T478K, E484A (present in most Omicron subvariants) and F486V (present in BA.4/5 and BQ.1.1) are well tolerated, suggesting some level of plasticity in the mode of antibody binding. BQ.1.1 has been shown to be resistant to neutralization by all approved therapeutic mAbs, including bebtelovimab ^1,12^, and became prevalent in the population worldwide, probably due to its advantage in evading humoral responses ^33^. Other new Omicron sublineages, such as XBB.1.5, a BA.2-derived recombinant, have been recently identified in the United States ^34^. This variant differs from BA.2 at ten residues of the RBD; however, none of the mutations affect the central 17T2 binding site, but rather they are located in the outer contact area and involve residues mutated in 17T2 sensitive variants (new mutation F486P and reversion R493Q)^5^.

All things considered, novel broadly-active neutralizing mAbs with proven *in vivo* neutralizing activity, such as 17T2 mAb, are urgently needed, especially to treat immunocompromised patients and those individuals at high risk of developing severe COVID-19. While only a few human mAbs have shown resilience against Omicron sublineages, antibodies SA58 and SA55 have shown partial or full coverage in the Omicron landscape ^5^. Interestingly, these antibodies bind to the RBD in a region distant from the 17T2 epitope, and structural data^35^ suggests that the RBD could simultaneously accommodate all three broadly neutralizing antibodies. Therefore, the combined use of these antibodies could improve their clinical efficacy, as previously suggested for combinations of mAbs active against pre-Omicron variants ^36^.

The identification of an anti-RBM broadly neutralizing antibody has several implications for future COVID-19 pandemic management. First, 17T2 has been cloned from a circulating IgA^+^ memory B cell isolated from a convalescent COVID-19 patient infected with the ancestral SARS-CoV-2. Therefore, our study provides evidence that infection with the ancestral virus could elicit broadly neutralizing antibodies, likely of mucosal origin, to SARS-CoV-2 variants not yet circulating. Additionally, and considering the benefits afforded by antibody-based therapies to treat COVID-19 patients and the excellent and broad neutralizing activity of 17T2 mAb *in vitro* and *in vivo*, we believe that 17T2 represents a promising candidate for future therapeutic and prophylactic interventions, either alone or in combination with other antibodies.

## Material and Methods

### Human subjects and sample collection

Blood samples were collected from a COVID-19 convalescent individual infected with SARS-CoV-2 during the first wave of the COVID-19 pandemic in Spain as previously described ^13^. Diagnosis of SARS-CoV-2 infection was confirmed by reverse transcription–quantitative polymerase chain reaction (RT-qPCR) of nasopharyngeal swab. Peripheral blood mononuclear cells (PBMCs) were isolated from whole blood collected with EDTA anticoagulant via Ficoll–Paque Premium (Cytiva, Freiburg, Germany) following the manufacturer’s instructions. PBMCs were resuspended in fetal bovine serum (FBS; Gibco, Riverside, MA, USA) with 10% dimethyl sulphoxide (DMSO; Sigma-Aldrich, St. Louis, USA) and stored in liquid nitrogen prior to use. All procedures were approved by the Ethical Committee for Clinical Investigation of the Institut Hospital del Mar d’Investigacions Mèdiques (Number 2020/9189/I).

### Single B-cell FACS sorting

For the isolation of WH1 SARS-CoV-2 spike-specific B cells, 26.4 pmol His-Tagged Biotinylated SARS-CoV-2 (2019-nCoV) spike RBD Recombinant Protein (Sino Biological Inc.; cat number: 40592-VO8H-B) was firstly incubated for 1 hour with 3.78 pmol Streptavidin Alexa Fluor 647 (Thermo Fisher Scientific) and Streptavidin Alexa Fluor 488 (Thermo Fisher Scientific), separately. Next, PBMCs were incubated with the fluorescently labelled RBD probes and with a cocktail of fluorescent conjugated antibodies containing anti-CD19 Pe-Cy7, anti-IgM BV605, anti HLA-DR AF700, anti CD38 APC-cy7, anti Igλ light chain PerCP-Cy5.5 (all from Biolegend), anti-IgD PE CF594 (BD Bioscience), anti-CD21 PE-cy5 (BD Bioscience) and anti-IgA Viogreen (Miltenyi). Dead cells were excluded through the use of 4’-6’-diamidine-2′-phenylindole (DAPI) (Sigma). Alive DAPI^-^ CD19^+^RBD^+^ B cells were single-cell index sorted using a FACSAria II (BD Biosciences) into empty 96-well PCR plates (VWR). FACSDiva software (Becton Dickinson) was used for acquisition and FlowJo for post-sorting analysis. Plates were immediately sealed with foil, frozen on dry ice and stored at −80 °C.

### Expression-cloning of antibodies

Antibodies were identified and sequenced as previously described ^14^. In brief, RNA from single cells was reverse transcribed in the original 96-well sort plate using random hexamers (Thermo Fisher Scientific), 0.76% NP 40 detergent solution (Thermo Fisher-Scientific), RNasin ribonuclease inhibitor (Promega), DTT (Invitrogen) and Superscript III Reverse Transcriptase (Invitrogen, 180080-044). The resulting cDNA was stored at −20 °C for subsequent amplification of the variable IGH, IGL and IGK genes by nested PCR and Sanger sequencing using same reverse and forward primers as in ^14^. Sequences were analyzed using Change-O (IgBlast). Following analysis, Ig gene-specific PCR was performed for the successfully annotated transcripts. Amplicon from the first PCR reaction were used as templates for cloning into antibody expression vectors (Abvec2.0-IGHG1 for the heavy chain and Abvec1.1-IGKC or Abvec1.1.-IGLC2-XhoI for of the light chains, all from AddGene).

### Production of the mAbs and Fabs

Both plasmids encoding for the LC and HC were transfected into Expi293F human cells (Thermo Fisher Scientific) with purified DNA and polyethylenimine (PEI 1:3). Cells were harvested by centrifugation at 4000 rpm for 5min after 5 days of expression at 37°C, 5% CO_2_ and 80% humidity. The cell supernatant containing the secreted mAb was purified by affinity chromatography in a HiTrap MabSelect (Cytiva) equilibrated in PBS and eluted with 10 mM Glycine at pH 3.

The purified 17T2 mAb was digested by incubation at 37°C within the immobilized papain agarose resin according to the manufacturer’s instructions (Thermo Fisher Scientific, No. 20341). The fragment antigen-binding (Fab) part was separated from undigested IgG and the crystallizable fragment (Fc) using an Immobilized Protein A column (Thermo Fisher Scientific, No. 20356). The flow through containing the Fab was concentrated through 10 kDa Amicon centrifugal filter units (EMD Millipore) and buffer exchange was performed with a 7 kDa molecular weight cut-off size exclusion resin (Thermo Fisher Scientific) to cryo-EM buffer (10 mM Tris at pH 7.6 and 20 mM NaCl).

### Production of recombinant SARS-CoV-2 proteins

The pCAGGS RBD construct, encoding for the RBD of the WH1 SARS-CoV-2 spike protein from the earliest lineage A virus (WH1, YP_009724390.1, residues 319-541; NC_045512.2, A lineage), along with the signal peptide plus a hexahistidine tag, was provided by Dr. Krammer (Mount Sinai School of Medicine, NY USA). RBD sequences from current Alpha, Beta, and Gamma VOCs were obtained from the World Health Organization tracking of variants (https://www.who.int/en/activities/tracking-SARS-CoV-2-variants/) and Pango lineage classification (https://cov-lineages.org/). DNA fragments encoding the RBD from Alpha (first identified in United Kingdom, B.1.1.7: N501Y), Beta (first identified in South Africa, B.1.351: K417N, E484K, N501Y) and Gamma (first identified in Japan/Brazil, P.1: K417T, E484K, N501Y) variants were synthetized by integrated DNA technology (IDT) as gblocks and codon optimized for mammalian expression. The fragments were inserted in a pCAGGS vector using Gibson Assembly. RBD proteins were expressed in-house in Expi293F human cells (Thermo Fisher Scientific) by transfection of the cells with purified DNA and polyethylenimine (PEI). RBD from Delta variant (first identified in India, B.1.617.2: L452R, T478K), Omicron and Omicron BA.2 were purchased from Sino Biological.

### ELISAs

96-well half-area flat bottom high-bind microplates (Corning) were coated overnight at 4°C with the different SARS-CoV-2 RBD recombinant viral proteins at 2 μg/mL in PBS (30 μl per well). Plates were washed with PBS 0.05% Tween 20 (PBS-T) and blocked with blocking buffer (PBS containing 1.5% Bovine serum albumin, BSA) for 2 hours at room temperature (RT). Monoclonal antibodies were serially diluted (starting dilution 10 μg/mL and then 11 serial dilutions 1:4) in PBS supplemented with 0.05% Tween 20 and 1% BSA and added to the viral protein-or PBS-coated plates for 2 hours at RT. After washing, plates were incubated with horseradish peroxidase (HRP)-conjugated anti-human IgG secondary antibody (Southern Biotech, 2042-05) diluted 1:4000 in PBS containing 0.05% Tween 20 and 1% BSA for 45 min at RT. Human IgG1 purified from serum of a myeloma patient (binding site company, BP078) was used as a negative control. Plates were washed 5 times with PBS-T and developed with TMB substrate reagent set (BD bioscience) with development reaction stopped with 1 M H_2_SO_4_. Absorbance was measured at 450 nm on a microplate reader (Infinite 200 PRO, Tecan). Optical density (OD) measurement was obtained after subtracting the absorbance at 570 nm from the absorbance at 450 nm.

### Pseudovirus generation and neutralization assay

HIV reporter pseudoviruses expressing SARS-CoV-2 spike protein and Luciferase were generated as previously described ^37^. pNL4-3.Luc.R-E-vector was obtained from the NIH AIDS Reagent Program ^38^. SARS-CoV-2.SctΔ19 was generated (GeneArt) from the full protein sequence of the ancestral SARS-CoV-2 spike (UniPro.org: P0DTC2) with a deletion of the last 19 amino acids in C-terminal ^39^, human-codon optimized and inserted into pcDNA3.1 (+). A similar procedure was followed to generate expression plasmids for all the different variants of SARS-CoV-2 spike according to consensus data (www.outbreak.info/) (**Supplementary Table 3**) as well as SARS-CoV spike (UniPro.org: P59594). Expi293F cells were transfected using ExpiFectamine293 Reagent (Thermo Fisher Scientific) with pNL4-3.Luc.R-.E- and SARS-CoV-2.SctΔ19 at an 8:1 ratio, respectively. Control pseudoviruses were obtained by replacing the spike protein expression plasmid with a VSV-G protein expression plasmid. Supernatants were harvested 48 hours after transfection, filtered at 0.45 mm, frozen, and titrated on HEK293T cells overexpressing wild-type human ACE-2 (Integral Molecular).

The SARS-CoV-2 pseudovirus-based neutralization assay were performed in Nunc 96-well cell culture plates (Thermo Fisher Scientific). Briefly, 200 TCID50 of pseudovirus were preincubated with purified monoclonal antibodies at 1 μg/ml and 1:5 dilution in PBS for 1 hour at 37°C. Then, 1×10^4^ HEK293T/hACE2 cells treated with DEAE-Dextran (Sigma-Aldrich) were added. Results were read after 48 hours using the EnSight Multimode Plate Reader and BriteLite Plus Luciferase reagent (PerkinElmer). The values were normalized and the half-maximal inhibitory concentrations (IC50) of the evaluated antibody were determined by plotting and fitting the log of the antibody concentration versus response to a 4-parameters equation in GraphPad Prism 9.0.0 (GraphPad Software).

### SARS-CoV-2 infection and prophylactic antibody treatment in K18-hACE2 mice

All animal procedures were performed under the approval of the Committee on the Ethics of Animal Experimentation of the IGTP and the authorization of Generalitat de Catalunya (code: 11222). B6. Cg-Tg(K18-ACE2)2Prlmn/J (or K18-hACE2) hemizygous transgenic mice (034860, Jackson Immunoresearch, West Grove, PA, United States) were bred, genotyped and maintained at CMCiB as stated in ^26^. A total of 24 adult K18-hACE2 hemizygous mice (aged 6-14 weeks) were used in this experiment, distributed in sex-balanced groups. One day before SARS-CoV-2 challenge, mice were anesthetized with isoflurane (FDG9623; Baxter, Deerfield, IL, USA) and administered by intraperitoneal injection with 10 mg/kg of either 17T2 mAb (n = 10) or and isotype control (anti-HIV-1 IgGb12) (n = 10).

After 24 hours, treated animals were challenged with 1000 TCID_50_ of Omicron BA.1.1 SARS-CoV-2 isolate (EPI_ISL_8151031), or PBS (uninfected control group). Viral challenge was performed under isoflurane anesthesia in 50 μL PBS (25 μL/nostril). All mice fully recovered from the infection and anesthesia procedures. Body weight and clinical signs were monitored daily from the antibody injection until the end of the experiment.

Five animals per treated group and two per uninfected group were euthanized 3 and 7 dpi for viral RNA quantification and pathological analyses. No animals had to be euthanized due to Humane Endpoint Criteria considering body weight loss and clinical signs ^26^. Euthanasia was performed under deep isoflurane anesthesia by whole blood extraction via intracardiac puncture followed by cervical dislocation. Oropharyngeal swab, lung, and nasal turbinate were collected for cell free viral RNA quantification. Lung tissue was collected for histological and immunohistochemistry analysis. SARS-CoV-2 PCR detection and viral load quantification of the samples was performed as described in ^26^.

### Histopathology and SARS-CoV-2 immunohistochemistry

Lung from mice were collected on days 3 and 7 post-SARS-CoV-2 inoculation, fixed by immersion in 10% buffered formalin and embedded into paraffin blocks. The histopathological analysis was performed on slides stained with hematoxylin/eosin and examined by optical microscopy. A semi-quantitative score based on the level of broncho-interstitial pneumonia (0-No lesion; 1-Mild, 2-Moderate or 3-Severe lesion) was established based on previous classifications ^40,41^. SARS-CoV-2 nucleoprotein was detected by immunohistochemistry (IHC) using the rabbit monoclonal antibody (40143-R019, Sino Biological) at a 1:15000 dilution. For immunolabelling visualization, the EnVision®+ System linked to horseradish peroxidase (HRP, Agilent-Dako) and 3,3’-diaminobenzidine (DAB) were used. The amount of viral antigen in tissues was semi-quantitatively scored (0-No antigen detection, 1-Low, 2-Moderate and 3-High amount of antigen) following previously published classifications ^40,41^.

### Determination of binding kinetics by surface plasmon resonance

Binding kinetics and affinity of 17T2 mAb for RBD were evaluated by surface plasmon resonance on a BIAcore T100 instrument (Cytiva) with a running buffer composed of 10 mM HEPES at pH 7.2, 150 mM NaCl and 0.05% Tween 20. The assay format involved antibody capture on a Series S CM5 chip. Briefly, amine coupling was used to create a human IgG capture surface (anti-human Fc mAb) following instructions provided with the Cytiva human IgG capture kit. 17T2 mAb was captured on flow cell 2, leaving flow cell 1 as a subtractive reference. Capture levels of IgG were targeted between 100 and 200 resonance units, after which serial dilution of RBD were flowed over immobilized IgG (50 μl/min for 2 min) and allowed to dissociate up to 30 min. The capture surface was regenerated with a 60-s injection of 3M MgCl_2_ (50 μl/min for 1 min). A 2-fold concentration series of each RBD variants ranging from 2 to 0,25 nM was used to analyze binding to 17T2. All sensograms were analyzed using a 1:1 Langmuir binding model with software supplied by the manufacturer to calculate the kinetics and binding constants. Where no decay in the binding signal was observed during the time allowed for dissociation, K_D_ based on the kd limit was determined by the ‘‘5% rule ^42^.

### De-glycosylation of monoclonal antibodies

Following the manufacturer’s instructions for PNGase F (Promega Inc., V483A) treatment, 200 μg of 17T2 mAb in 50 mM ammonium bicarbornate (pH 7.4) was combined with 50 μl of Glyco Buffer 10X and water to make up a 500 μl total reaction volume. The mixture was incubated at 37°C overnight without or with 20 μl of PNGase F. Control analysis of the mAb de-glycosylation was performed by gel-shift on SDS-PAGE. The mAb preparation was concentrated through a 50 kDa Amicon centrifugal filter to remove the PNGase enzyme (EMD Millipore) and dialyzed in PBS 1X for further experiments.

### Cryo-electron microscopy sample preparation and data acquisition

B.1.1.529 (BA.1) S1 + S2 Trimer-His Recombinant Protein (Sino Biologicals; 40589-V08H26) was reconstituted in sterile water (100 μl) to prepare a stock solution (1mg/ml). Buffer exchange to 10 mM Tris pH 7.6 was performed twice through ZebaTM Spin Desalting Columns (Thermo Scientific, number 899882). BA.1 spike was mixed with the Fab 17T2 (1:1.5 molar ratio of spike monomer: Fab, i.e. 0.59:0.71 mg/mL, respectively) and 3 μl were kept 5 min at RT until their application onto glow-discharged holey carbon grids (Quantifoil, Au 300 mesh, R 0.6/1). The grids were blotted and then plunged into liquid ethane using a FEI Vitrobot Mark IV at 20°C and 95% relative humidity. Data were collected at a FEI Talos Arctica electron microscope operated at 200 kV and equipped with a Falcon III electron detector. A total of 9,615 movies (**Supplementary Figure 3A**) were recorded at a defocus range of −1 μm to −2.5 μm with a pixel size of 0.855 Å; exposure time was 40 s, with a total exposure dose of 32 e/Å2 over 60 frames.

### Image processing

All image processing steps were performed inside Scipion ^43^. We used Scipion 3.0 in order to easily combine different software suites in the analysis workflows of cryo-EM data: movie frames were aligned using Relion’s implementation of the UCSF MotionCor2 program ^44,45^. The contrast transfer function (CTF) of the micrographs was estimated using GCTF ^46^. Movies were then automatically picked using Gautomatch. Following the application of the Scipion picking consensus protocol, 1,203,207 particles were extracted. 2D classification was performed in cryoSPARC ^47^ and 124,570 particles were selected (**Supplementary Figure 3B**). The cryoSPARC initial model protocol was then used to generate and classify the particles into 2 classes, without imposing symmetry (**Supplementary Figure 3C**). Class 1 contained low quality particles (16,866) and class 2 contained a dataset with the highest number of particles with high quality particles (107,704 particles). The highest dataset was selected to perform a further classification yielding indistinguishable classes in which all the RBDs were in up conformation (Data not shown). The dataset with the highest number of particles (107,704 particles) was refined using non-uniform refinement in cryoSPARC with no symmetry application (**Supplementary Figure 3D**), to overall resolution of 3.46 Å based on the gold-standard (FSC = 0.143) criterion (**Supplementary Figure 3F**). The resulting map was sharpened with DeepEMhancer ^48^.

### Model building and refinement

The coordinates of the SARS-CoV-2 spike in PDB ID: 7Y9S were used as initial model for fitting the cryo-EM map. Output from AlphaFold 2.0 modelling was used as initial model for Fab 17T2. Due to the high flexibility in RBD-antibody region and consequently the lower resolution in this area, local refinements with CryoSPARC were performed, using a mask encompassing the RBD-Fab region (**Supplementary Figure 3E**). The final resolution was 4.41 FÅ map, which allowed for increased definition in this region (**Supplementary Figure 3G**). However, the side chains were not fully resolved. Iterative manual model building and real-space refinement were carried out in Coot ^49^ and in Phenix ^50^ and CCP-EM^51^. The validation of the model was done with Molprobity ^52^ (**Supplementary Table 5**), sofware integrated in Phenix suite. UCSF Chimera and ChimeraX were used for map fitting and manipulation ^53^.

### Statistical analysis

All figures were generated in GraphPad Prism 9.0.0. Statistical analyses were performed using R v4.1.1. Unpaired datasets were analyzed using a Kruskal-Wallis with Dunn’s correction for multiple testing. Histopathological and IHC scores were compared using the Generalized Pearson Chi-Squared Test for ordinal data. Viral load comparisons were analyzed using a Petro-Prentice generalized Wilcoxon test.

## Acknowledgements

We acknowledge access to the cryo-EM CNB-CSIC facility in the context of the CRIOMECORR project (ESFRI-2019-01-CSIC-16) and we thank the staff of the Protein Technology Unity (CRG) for the help in protein production. This study was supported by the COVID-19 call grant from Generalitat de Catalunya, Department of Health (to GM), grant Miguel Servet research program (to GM), and partially funded by the crowdfunding initiative #joemcorono and the Fundació Glòria Soler (to JB). A.P-G. was supported by a predoctoral grant from Generalitat de Catalunya and Fons Social Europeu (2022 FI_B 00698).

## Author Contributions

L.D.C.-M. produced the monoclonal antibodies, designed and performed experiments, analyzed and discussed data and reviewed and edited the manuscript. S.T.V. produced the monoclonal antibodies, designed and performed experiments and discussed data. N.R.M. performed SARS-CoV-2 antigens and mAbs expression and purification and carried out SPR experiments. B.T. carried out neutralization assays and *in vivo* experiments with K18-hACE-2 mice, analyzed and interpreted the data and contributed to writing of the manuscript. E.P. and S.M. carried out neutralization assays. A.P.-G., F.T.-F. and N.I. carried out *in vivo* experiments with K18-hACE-2 mice. E.R.-M. and E.B. performed VL quantification. J.V.-A. carried out immunohistochemistry and histology experiments and analyzed and interpreted the data. M.P. carried out immunohistochemistry and histology experiments. J.S. carried out immunohistochemistry and histology experiments and analyzed and interpreted the data. E.B. analyzed and interpreted the data. A.M. carried out computational aspects of image processing and structure determination, interpreted and analyzed the cryo-EM structures and binding analyses and carried out experimental aspects of cryo-EM. M.T.B.-C. collected cryo-EM data. D.C. carried out computational aspects of image processing and structure determination. C.S. interpreted and analyzed the cryo-EM structures and binding analyses. R.A. carried out experimental aspects of cryo-EM, interpreted and analyzed the cryo-EM structures and binding analyses and wrote the manuscript. C.C. designed and performed experiments, carried out SPR experiments, discussed data and reviewed and edited the manuscript. J.B analyzed and interpreted the data and wrote the manuscript. G.M. designed the study, produced the monoclonal antibodies, analyzed and interpreted data, and wrote the manuscript.

## Data and materials availability

The data that support this study are available from the corresponding authors upon reasonable request. Cryo-EM data have been deposited in the Electron Microscopy Data Bank under accession codes EMD-16453 for SARS-CoV-2 Spike Trimer in complex three 17T2 Fabs and EMD-1643 for RBD/17T2 Fab and its associated atomic models have been deposited in the Protein Data Bank under accession code 8C89. Further information and requests for resources and reagents should be directed to and will be fulfilled by the lead Contact, Giuliana Magri.

