## Supplementary Material for "A Novel Monoclonal Antibody Targeting a Large Surface of the Receptor Binding Motif Shows Pan-neutralizing SARS-CoV-2 Activity Including BQ.1.1 Variant"

Supplementary Figures and legends 1-5
Supplementary Tables 1-4

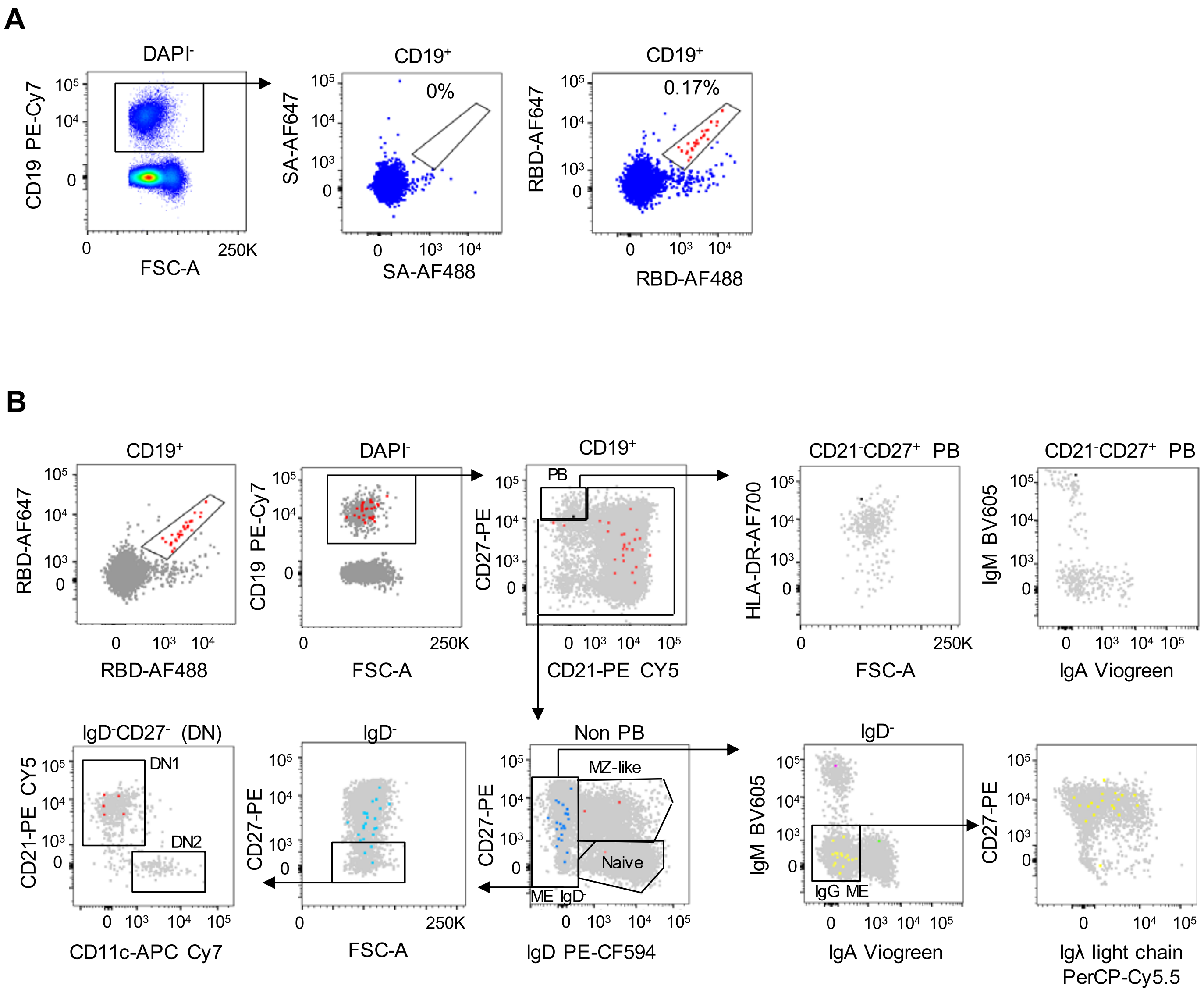
A
DAPI-
CD19 PE-Cy7
FSC-A
250K
CD19+
0%
SA-AF647
SA-AF488
CD19+
0.17%
RBD-AF647
RBD-AF488
B
CD19+
RBD-AF647
RBD-AF488
DAPI-
CD19 PE-Cy7
FSC-A
CD19+
PB
CD27-PE
CD21-PE CY5
CD21-CD27+ PB
HLA-DR-AF700
FSC-A
CD21-CD27+ PB
IgM BV605
IgA Viogreen
IgD-CD27- (DN)
DN1
DN2
CD21-PE CY5
CD11c-APC Cy7
IgD-
CD27-PE
FSC-A
Non PB
MZ-like
Naive
ME IgD-
CD27-PE
IgD PE-CF594
IgD-
IgM BV605
IgG ME
IgA Viogreen
CD27-PE
Igλ light chain
PerCP-Cy5.5

**Supplementary Figure 1. Characterization SARS CoV-2 WH1 RBD-specific B cells by flow cytometry**. **(A)** Flow cytometry staining of $CD19^{+}$ B cells from a convalescent COVID-19 individual without (left) and with (right) two fluorescently labeled biotinylated RBD probes. Numbers indicate the percentage of RBD-specific cells within total $CD19^{+}$ B cells. Red large dots represent cells that are positive for both RBD-AF647 and RBD-AF488. **(B)** Gating strategy used to define RBD-specific B cell populations: PBs ($CD19^{+}CD27^{++}CD21^{-}$), MZ-like (non-PB $CD19^{+}CD27^{+}IgD^{+}$), naïve (non-PB $CD19^{+}CD27^{-}IgD^{+}$), ME $IgD^{-}$ (non-PB $CD19^{+}IgD^{-}$), $IgM^{+}$ ME (non-PB $CD19^{+}IgD^{-}IgM^{+}$), $IgA^{+}$ ME (non-PB $CD19^{+}IgD^{-}IgA^{+}$), $IgG^{+}$ ME (non-PB $CD19^{+}IgD^{-}IgG^{+}$), DN1 (non-PB $CD19^{+}IgD^{-}CD27^{-}CD21^{+}CD11c^{-}$), and DN2 (non-PB $CD19^{+}IgD^{-}CD27^{-}CD21^{-}CD11c^{+}$). RBD-specific cells are represented in colored large dots.

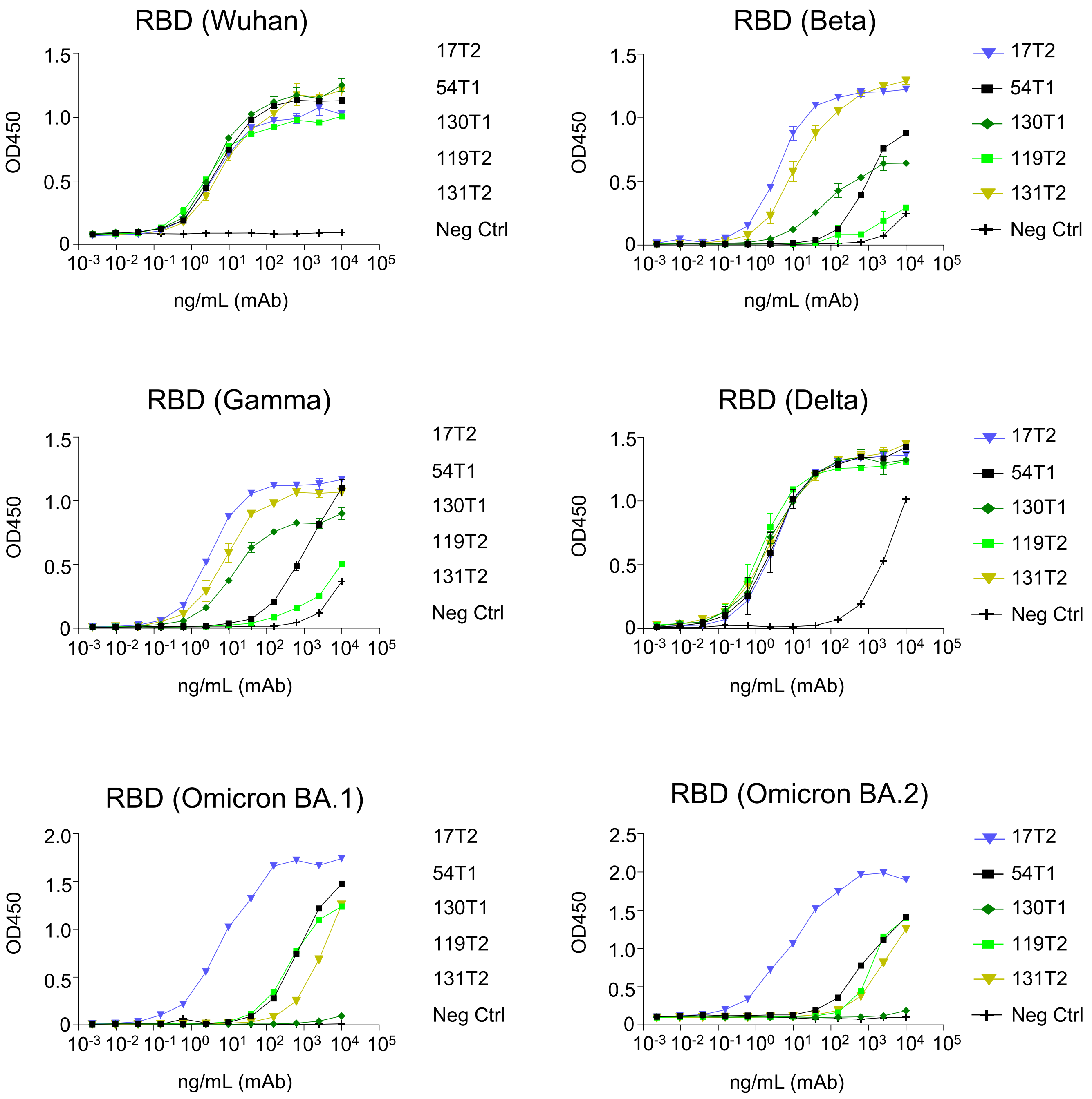
RBD (Wuhan)
RBD (Beta)
RBD (Gamma)
RBD (Delta)
RBD (Omicron BA.1)
RBD (Omicron BA.2)
OD450
ng/mL (mAb)
0
0.5
1.0
1.5
2.0
2.5
10-3 10-2 10-1 100 101 102 103 104 105
17T2
54T1
130T1
119T2
131T2
Neg Ctrl

**Supplementary Figure 2. Reactivity of SARS-CoV-2 RBD-specific monoclonal antibodies**. ELISA binding curves of serial dilutions of the mAbs to the spike RBD of SARS-CoV-2 from different variants, coated at equimolar concentrations. Graph bars represent the average ± SD. Human IgG1 purified from serum of a myeloma patient (binding site company, BP078) was used as a negative control for binding.

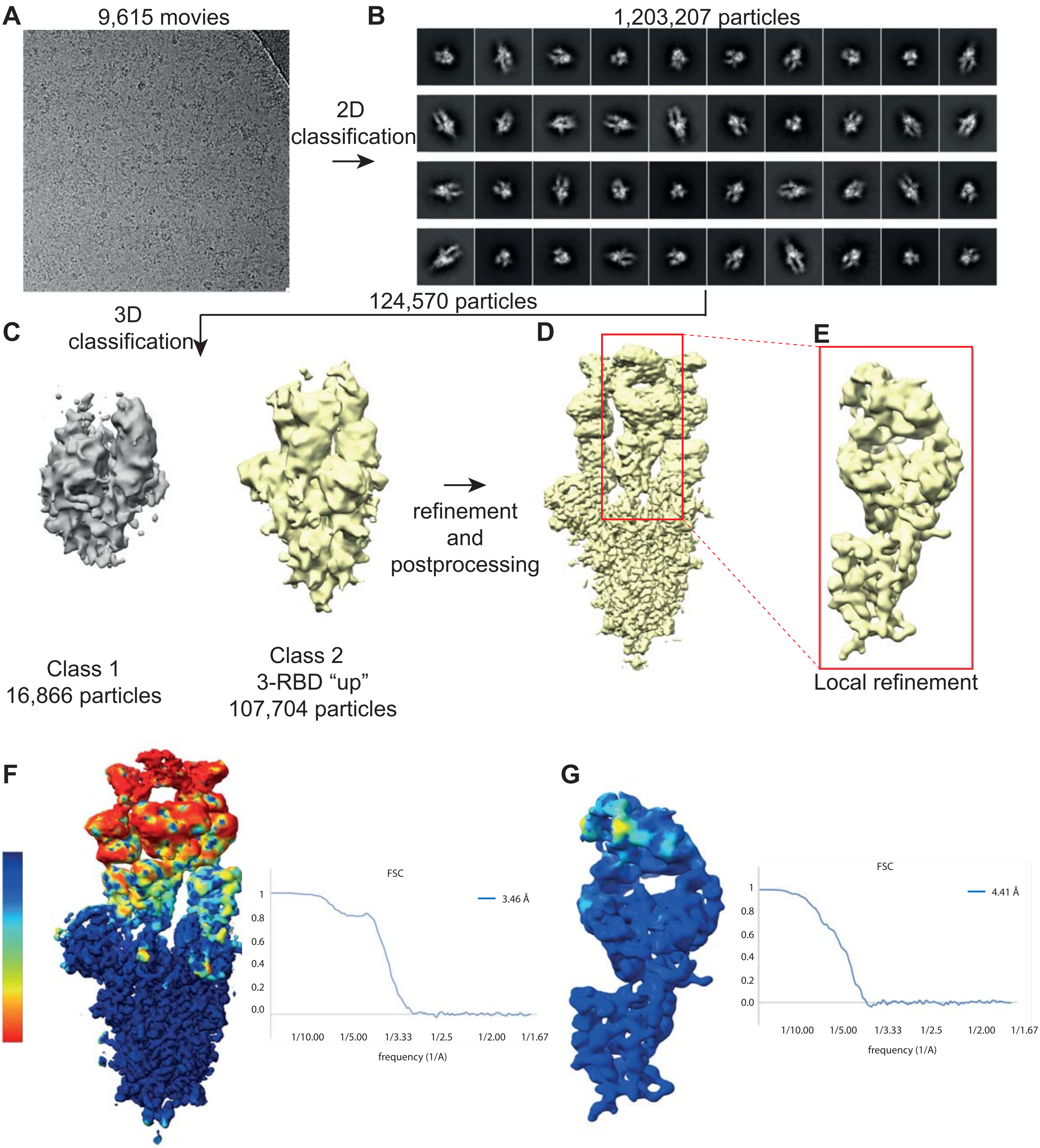

A
9,615 movies
2D
classification
B
1,203,207 particles
124,570 particles
C
3D
classification
Class 1
16,866 particles
Class 2
3-RBD “up”
107,704 particles
refinement
and
postprocessing
D
E
Local refinement
F
3.5
4.8
6.1
7.4
8.7
10.0
FSC
3.46 Å
1
0.8
0.6
0.4
0.2
0.0
1/10.00
1/5.00
1/3.33
1/2.5
1/2.00
1/1.67
frequency (1/A)
G
FSC
4.41 Å
1
0.8
0.6
0.4
0.2
0.0
1/10.00
1/5.00
1/3.33
1/2.5
1/2.00
1/1.67
frequency (1/A)

**Supplementary Figure 3. Cryo-EM data analysis for complex between the Omicron BA.1 spike trimer and the 17T2 Fab fragment.** **(A)** Representative electron micrograph. **(B)** Representative 2D-class averages. **(C)** 3D classification. Left class contains low quality particles and right class contains high quality particles that were selected for refinement. **(D)** 3D reconstruction of complex with no symmetric imposed. **(E)** The boxed region contains one RBD complexed with one Fab in a refined map masked by local refinement. **(F-G)** Gold standard Fourier shell correlation (FSC) curve of final overall (left) and locally refined (right) maps and resolution estimation based on 0.143 Fourier shell correlation criteria as indicated by a blue line.

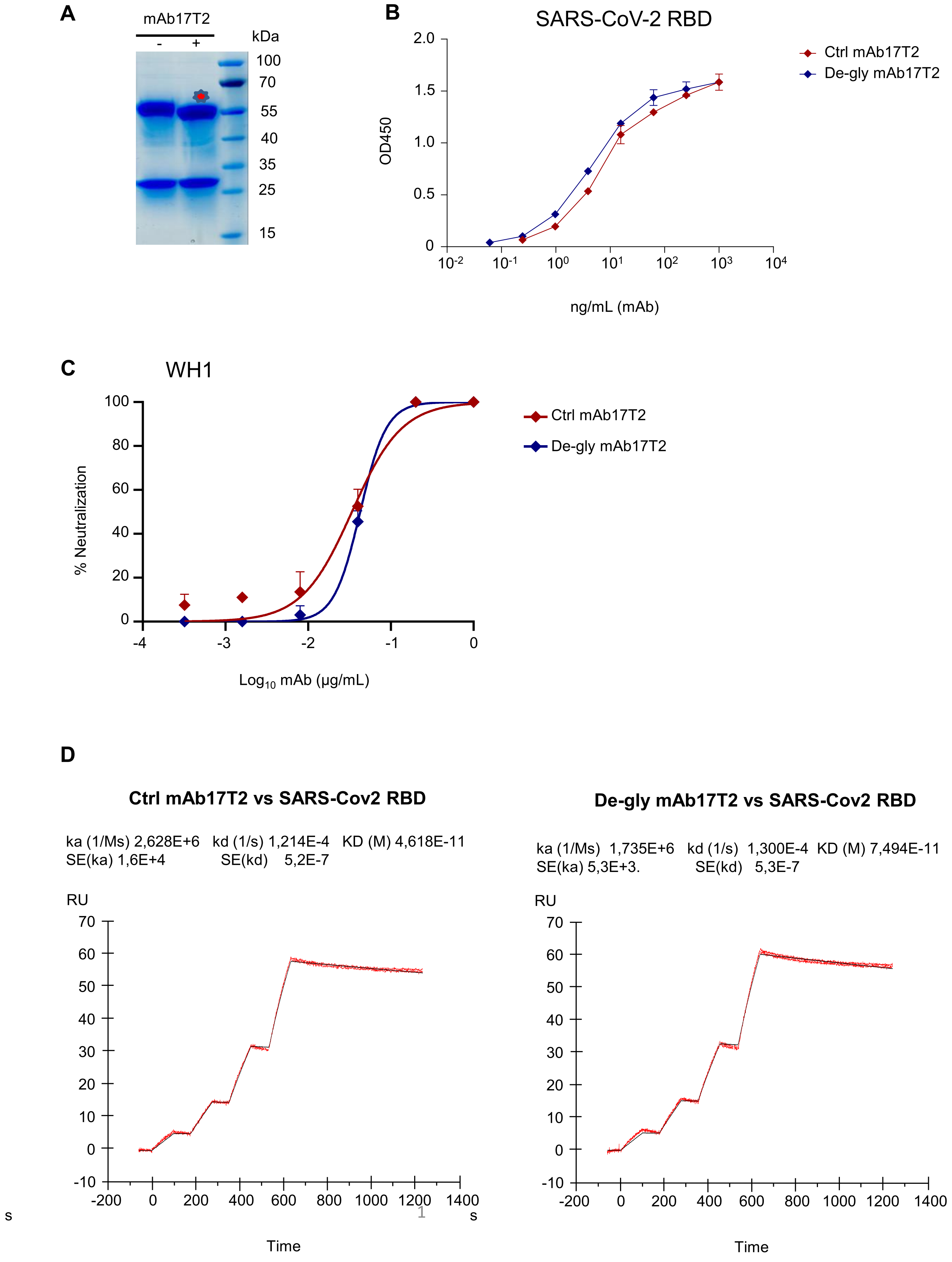
A
mAb17T2
- +
kDa
100
70
55
40
35
25
15
B
SARS-CoV-2 RBD
OD450
2.0
1.5
1.0
0.5
0
10-2 10-1 10⁰ 10¹ 10² 10³ 10⁴
ng/mL (mAb)
Ctrl mAb17T2
De-gly mAb17T2
C
WH1
% Neutralization
100
80
60
40
20
0
-4 -3 -2 -1 0
Log10 mAb (μg/mL)
Ctrl mAb17T2
De-gly mAb17T2
D
Ctrl mAb17T2 vs SARS-Cov2 RBD
ka (1/Ms) 2,628E+6 kd (1/s) 1,214E-4 KD (M) 4,618E-11
SE(ka) 1,6E+4 SE(kd) 5,2E-7
RU
70
60
50
40
30
20
10
0
-10
-200 0 200 400 600 800 1000 1200 1400
s
s
Time
De-gly mAb17T2 vs SARS-Cov2 RBD
ka (1/Ms) 1,735E+6 kd (1/s) 1,300E-4 KD (M) 7,494E-11
SE(ka) 5,3E+3. SE(kd) 5,3E-7
RU
70
60
50
40
30
20
10
0
-10
-200 0 200 400 600 800 1000 1200 1400
Time

**Supplementary Figure 4. PNGase F treatment of purified 17T2 mAb. (A)** Purified 17T2 mAb was incubated in the presence (+) or absence (–) of recombinant PNGase F and analyzed by SDS-PAGE. Red asterisks indicate the de-glycosylated product. **(B)** ELISA binding curves of serial dilutions of control treated 17T2 (red) and de-glycosylated 17T2 (blue) mAbs to recombinant SARS-CoV-2 RBD, coated at equimolar concentrations. Graph bars represent the average ± SD. **(C)** Neutralization curves of control treated 17T2 (red) and de-glycosylated 17T2 (blue) mAbs against ancestral SARS-CoV-2 WH1 variant. Values are mean +/- SD of duplicate samples. **(D)** The binding kinetics of de-glycosylated (left) and no de-glycosylated form of 17T2 mAb (right) to recombinant SARS-CoV-2 RBD were obtained using the BIAcore T100 system in single-cycle mode. mAbs were captured on the chip, and serial dilutions of RBD (WH1) were then injected over the chip surface. The $K_D$ is labelled accordingly.

**A**

Heavy chain

| | CDR H1 17T2 | | | | | | | |
|---|---|---|---|---|---|---|---|---|
| | 26 | 27 | 28 | 29 | 30 | 31 | 32 | 33 |
| 17T2 | V | F | T | F | S | I | S | A |
| S2E12 | G | F | T | F | T | S | S | A |

| | CDR H2 17T2 | | | | | | | |
|---|---|---|---|---|---|---|---|---|
| | 51 | 52 | 53 | 54 | 55 | 56 | 57 | 58 |
| 17T2 | I | V | V | G | S | G | N | T |
| S2E12 | I | V | V | G | S | G | N | T |

| | CDR H3 17T2 | | | | | | | | | | | | | | | |
|---|---|---|---|---|---|---|---|---|---|---|---|---|---|---|---|---|
| | 97 | 98 | 99 | 100 | 101 | 102 | 103 | 104 | 105 | 106 | 107 | 108 | 109 | 110 | 111 | 112 |
| 17T2 | A | A | P | Y | C | N | R | T | T | C | Y | D | G | F | D | L |
| S2E12 | A | S | P | Y | C | S | G | G | S | C | S | D | G | F | D | I |

Light chain

| | CDR L1 17T2 | | | | | | |
|---|---|---|---|---|---|---|---|
| | 27 | 28 | 29 | 30 | 31 | 32 | 33 |
| 17T2 | Q | S | I | S | S | N | Y |
| S2E12 | Q | S | V | S | S | S | Y |

| | CDR L2 17T2 | | |
|---|---|---|---|
| | 51 | 52 | 53 |
| 17T2 | G | A | S |
| S2E12 | G | A | S |

| | CDR L3 17T2 | | | | | | | | | |
|---|---|---|---|---|---|---|---|---|---|---|
| | 90 | 91 | 92 | 93 | 94 | 95 | 96 | 97 | 98 | 99 |
| 17T2 | Q | H | Y | G | G | L | S | R | W | T |
| S2E12 | Q | Q | Y | V | G | L | T | G | W | T |

**B**

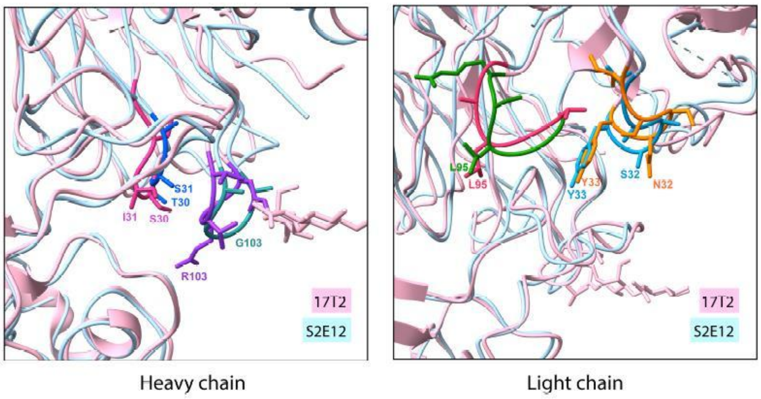


**C**

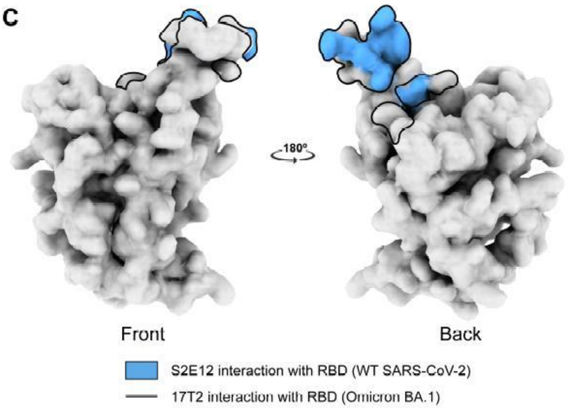

**Supplementary Figure 5. Comparative study of 17T2 Fab with similar antibody S2E12 Fab.**

**(A)** Comparison of the sequences of the CDRs of both the heavy and light chains between the 17T2 and S2E12 antibodies, showing differences between residues with colors. **(B)** Alignment of the RBD-Fab structures of 17T2 and 2SE12 Fabs showing the side chains of the residues that are part of the regions with sequence differences (color codes as in **(A)**). **(C)** Different areas of interaction of 17T2 and S2E12 Fabs with the RBD of SARS-CoV-2 spike.

**Supplementary Table 1. Phenotype and native isotype of the B cells originating the mAbs, and the antibody gene usage of the heavy and light chain variable regions.**

| Name | B cell of origin | |
|---|---|---|
| | B cell phenotype | Isotype |
| **17T2** | CD19$^+$HLA-DR$^+$CD27$^+$IgA$^+$CD21$^+$IgM$^-$IgD$^-$lambda$^-$ | IgA |
| **54T1** | CD19$^+$HLA-DR$^+$CD27$^+$IgA$^+$CD21$^+$IgM$^-$IgD$^-$lambda$^-$ | IgA |
| **130T1** | CD19$^+$HLA-DR$^+$CD27$^+$IgA$^-$CD21$^+$IgM$^-$IgD$^-$lambda$^-$ | IgG |
| **119T2** | CD19$^+$HLA-DR$^+$CD27$^+$IgA$^-$CD21$^+$IgM$^-$IgD$^-$lambda$^+$ | IgG |
| **131T2** | CD19$^+$HLA-DR$^+$CD27$^+$IgA$^-$CD21$^+$IgM$^-$IgD$^-$lambda$^-$ | IgG |

| Name | IgHeavy | | | |
|---|---|---|---|---|
| | V call | D call | J call | % V identity |
| **17T2** | IGHV1-58*01 | IGHD2-2*01,IGHD2-2*02,IGHD2-2*03 | IGHJ3*01,IGHJ3*02 | 94,5 |
| **54T1** | IGHV3-53*04 | IGHD5-12*01 | IGHJ6*02 | 96,6 |
| **130T1** | IGHV1-46*01 | IGHD5-18*01,IGHD5-5*01 | IGHJ4*01,IGHJ4*02 | 96,6 |
| **119T2** | IGHV3-9*01 | IGHD2-15*01,IGHD2-21*01,IGHD2-21*02 | IGHJ3*02 | 98 |
| **131T2** | IGHV3-30*18,IGHV | IGHD5-18*01,IGHD5-5*01 | IGHJ4*02 | 97,3 |

| Name | IgLight | | | |
|---|---|---|---|---|
| | Chain | V call | J call | % V identity |
| **17T2** | **Kappa** | IGKV3-20*01 | IGKJ1*01 | 96,5 |
| **54T1** | **Kappa** | IGKV1-9*01 | IGKJ1*01 | 98,6 |
| **130T1** | **Kappa** | IGKV1-17*01 | IGKJ1*01 | 98,6 |
| **119T2** | **Lambda** | IGLV2-14*03 | IGLJ3*02 | 98,3 |
| **131T2** | **Kappa** | IGKV3-15*01 | IGKJ4*01 | 97,6 |

**Supplementary Table 2. Kinetic rate constants and affinities determined for the 17T2 mAb against SARS-CoV-2 RBD variants.**

| 17T2 | | | |
|---|---|---|---|
| **SARS-CoV-2 RBD variants** | **ka (1/Ms)** | **kd (1/s)[a]** | **$K_D$ (nM)[b]** |
| WH1 | 2,62E+06 | 1,21E-4 | 0,0462 |
| Alpha | 9,61E+05 | <1.0E-05 | <0.1 |
| Beta | 2,30E+06 | <1.0E-05 | <0.1 |
| Delta | 3,32E+06 | <1.0E-05 | <0.1 |
| Omicron BA.1 | 1,08E+07 | <1.0E-05 | <0.1 |
| Omicron BA.2 | 5,90E+07 | <1.0E-05 | <0.1 |

a. For most of the Biacore experiments, no decay in the binding signal was observed during the time allowed for dissociation. According to the ‘‘5% rule’’, the $k_d$ can be resolved only if 5% or more of the bound material dissociates. Therefore, the upper limit on the $K_d$ (in 1/s) is given by $K_d < \ln(0.95)/td$, where td is the amount of time (in seconds) allowed for dissociation.

b. $K_D$ based on the kd limit determined by the ‘‘5% rule.’’

**Supplementary Table 3. Pseudovirus spike mutations relative to ancestral WH1.**

| Name | Mutations |
|---|---|
| D614G | D614G |
| Alpha | H69-70del, Y144del, N501Y, A570D, P681H, T716I, S982A and D1118H |
| Beta | L18F, D80S, D215G, L242-244del, R246I, K417N, E484K, N501Y, D614G, A701V |
| Gamma | L18F, T20N, P26S, D138Y, R190S, K417T, E484K, N501Y, D614G, H655Y, T1027I, V1176F |
| Delta | T19R, 157-158 del, L452R, T478K, D614G, P681R, D950N |
| Mu | T95I, Y144S, Y145N, R346K, E484K, N501Y, D614G, P681H, D950N |
| BA.1 | A67V, H69-70del, T95I, G142D, V143-145del, N211del, L212I, ins214EPE, G339D, S371L, S373P, S375F, K417N, N440K, G446S, S477N, T478K, E484A, Q493R, G496S, Q498R, N501Y, Y505H, T547K, D614G, H655Y, N679K, P681H, N764K, D796Y, N856K, Q954H, N969K, L981F |
| BA.2 | T19I, L24S, P25-27del, G142D, V213G, G339D, S371F, S373P, S375F, T376A, D405N, R408S, K417N, N440K, S477N, T478K, E484A, Q493R, Q498R, N501Y, Y505H, D614G, H655Y, N679K, P681H, N764K, D796Y, Q954H, N969K |
| BA.4/5 | T19I, L24S, P25-27del, H69-70del, G142D, V213G, G339D, S371F, S373P, S375F, T376A, D405N, R408S, K417N, N440K, L452R, S477N, T478K, E484A, F486V, Q498R, N501Y, Y505H, D614G, H655Y, N679K, P681H, N764K, D796Y, Q954H, N969K |
| BQ.1.1 | T19I, L24S, P25-27del, H69-70del, G142D, V213G, G339D, R346T, S371F, S373P, S375F, T376A, D405N, R408S, K417N, N440K, K444T, L452R, N460K, S477N, T478K, E484A, F486V, Q498R, N501Y, Y505H, D614G, H655Y, N679K, P681H, N764K, D796Y, Q954H, N969K |

**Supplementary Table 4. Representation of reported SARS-CoV-2 RBD mutations in key variants.** Color highlighted boxes show the presence of RBD mutations in each SARS-CoV-2 variant (same color codes as in **Figure 3F).**

| | | Receptor-binding-domain mutations | | | | | | | | | | | | | | | | | | | | | | | |
|---|---|---|---|---|---|---|---|---|---|---|---|---|---|---|---|---|---|---|---|---|---|---|---|---|---|
| | | 339 | 346 | 371 | 373 | 375 | 376 | 405 | 408 | 417 | 440 | 444 | 446 | 452 | 460 | 477 | 478 | 484 | 486 | 490 | 493 | 496 | 498 | 501 | 505 |
| WT Sars-Cov-2 | | G | R | S | S | S | T | D | R | K | N | K | G | L | N | S | T | E | F | F | Q | G | Q | N | Y |
| Alpha | B.1.1.7 | | | | | | | | | | | | | | | | | | | | | | | Y | |
| Beta | B.1.351 | | | | | | | | | N | | | | | | | | K | | | | | | Y | |
| Gamma | P.1 | | | | | | | | | T | | | | | | | | K | | | | | | Y | |
| Delta | B.1.617.2 | | | | | | | | | | | | | R | | | K | | | | | | | | |
| Epsilon | B.1.427/9 | | | | | | | | | | | | | R | | | | | | | | | | | |
| Zeta | P.2 | | | | | | | | | | | | | | | | | K | | | | | | | |
| Eta | B.1.525 | | | | | | | | | | | | | | | | | K | | | | | | | |
| Theta | P.3 | | | | | | | | | | | | | | | | | K | | | | | | Y | |
| Iota | B.1.526 | | | | | | | | | | | | | | | | | K | | | | | | | |
| Kappa | B.1.617.1 | | | | | | | | | | | | | R | | | K | Q | | | | | | | |
| Lambda | C.37 | | | | | | | | | | | | | Q | | | | | | S | | | | | |
| Mu | B.1.621 | | K | | | | | | | | | | | | | | | K | | | | | | Y | |
| Omicron | BA.1 | D | | L | P | F | | | | N | K | | S | | | N | K | A | | | R | S | R | Y | H |
| | BA.2 | D | | F | P | F | A | N | S | N | K | | | | | N | K | A | | | R | | R | Y | H |
| | BA.4/BA.5 | D | | F | P | F | A | N | S | N | K | | | R | | N | K | A | V | | | | R | Y | H |
| | BQ.1.1 | D | | F | P | F | A | N | S | N | K | T | | R | K | N | K | A | V | | | | R | Y | H |

**Supplementary Table 5. RBD/17T2 Fab model refinement and statistic.**

| Model | RBD-Fab | |
|---|---|---|
| **Composition (#)** | | |
| **Chains** | 3 | |
| **Atoms** | 5106 (Hydrogens: 0) | |
| **Residues** | Protein: 652 Nucleotide: 0 | |
| **Water** | 0 | |
| **Ligands** | NAG: 4 | |
| **Bonds (RMSD)** | | |
| **Length (Å) (# > 4σ)** | 0.006 (0) | |
| **Angles (°) (# > 4σ)** | 1.208 (10) | |
| **MolProbity score** | 2.47 | |
| **Clash score** | 25.75 | |
| **Ramachandran plot (%)** | | |
| **Outliers** | 0.31 | |
| **Allowed** | 10.37 | |
| **Favored** | 89.32 | |
| **Rotamer outliers (%)** | 0.71 | |
| **Cβ outliers (%)** | 0.00 | |
| **Peptide plane (%)** | | |
| **Cis proline/general** | 5.4/0.2 | |
| **Twisted proline/general** | 0.0/0.0 | |
| **CaBLAM outliers (%)** | 5.00 | |
| **ADP (B-factors)** | | |
| **Iso/Aniso (#)** | 5106/0 | |
| **min/max/mean** | | |
| **Protein** | 23.54/440.00/243.00 | |
| **Nucleotide** | --- | |
| **Ligand** | 106.97/440.00/301.85 | |
| **Water** | --- | |
| **Occupancy** | | |
| **Mean** | 1.00 | |
| **occ = 1 (%)** | 100.00 | |
| **0 < occ < 1 (%)** | 0.00 | |
| **occ > 1 (%)** | 0.00 | |
| **Data** | | |
| **Box** | | |
| **Lengths (Å)** | 60.40, 66.15, 119.70 | |
| **Angles (°)** | 90.00, 90.00, 90.00 | |
| **Supplied Resolution (Å)** | 3.5 | |
| **Resolution Estimates (Å)** | Masked | Unmasked |
| **d FSC (half maps; 0.143)** | 3.6 | 3.8 |
| **d 99 (full/half1/half2)** | 2.4/5.5/5.5 | 2.4/5.1/5.1 |
| **d model** | 2.0 | 2.0 |
| **d FSC model (0/0.143/0.5)** | 1.9/2.6/3.5 | 1.9/2.6/3.5 |
| **Map min/max/mean** | -1.27/1.54/0.01 | |
| **Model vs. Data** | | |
| **CC (mask)** | 0.78 | |
| **CC (box)** | 0.79 | |
| **CC (peaks)** | 0.79 | |
| **CC (volume)** | 0.78 | |
| **Mean CC for ligands** | 0.84 | |
